# Comprehensive label-free characterization of extracellular vesicles and their surface proteins

**DOI:** 10.1101/2020.12.28.424566

**Authors:** E. Priglinger, J. Strasser, B. Buchroithner, F. Weber, S. Wolbank, D. Auer, E. Grasmann, C. Arzt, M-S. Narzt, J. Grillari, J. Preiner, J. Jacak, M. Gimona

## Abstract

Interest in mesenchymal stem cell derived extracellular vesicles (MSC-EVs) as therapeutic agents has dramatically increased over the last decade. Preclinical studies show that MSC-EVs have anti-apoptotic and neuroprotective effects, boost wound healing, and improve the integration of allogeneic grafts through immunomodulation. Current approaches to the characterization and quality control of EV-based therapeutics include particle tracking techniques, Western blotting, and advanced cytometry, but standardized methods are lacking. In this study, we established and verified quartz crystal microbalance (QCM) as highly sensitive label-free immunosensing technique for characterizing clinically approved umbilical cord MSC-EVs enriched by tangential flow filtration and ultracentrifugation. Using QCM in conjunction with common characterization methods, we were able to specifically detect EVs via EV (CD9, CD63, CD81) and MSC (CD44, CD49e, CD73) markers and gauge their prevalence. Additionally, we characterized the topography and elasticity of these EVs by atomic force microscopy (AFM), enabling us to distinguish between EVs and non-vesicular particles (NVPs) in a therapeutic formulation. This measurement modality makes it possible to identify EV sub-fractions, discriminate between EVs and NVPs, and to characterize EV surface proteins, all with minimal sample preparation and using label-free measurement devices with low barriers of entry for labs looking to widen their spectrum of characterization techniques. Our combination of QCM with impedance measurement (QCM-I) and AFM measurements provides a robust multi-marker approach to the characterization of clinically approved EV formulations and opens the door to improved quality control.

## Introduction

The interest in extracellular vesicles (EVs) for therapeutic use or as a medical product has increased massively over the past decade. EVs are membrane surrounded particles that are naturally released by potentially all eukaryotic cell types (Ludwig and Giebel, 2012). Exosomes or small EVs (sEVs) range in size from 30 to 150 nm and carry cell type-specific proteins, lipids and coding and non-coding ribonucleic acids. They have been implicated in cellular signaling to regulate biological processes such as immunomodulation and regeneration, but also play a pathophysiological role in cancer, infections, and degenerative diseases such as neurodegeneration (Rohde et al., 2019). Considering the above properties, EVs are promising new active biological agents that can be valuable for a wide variety of medical applications such as diagnostic biomarkers, vaccine carriers, drug delivery vectors for cytostatics in cancer therapy, and therapeutic agents for tissue regeneration (Melling et al., 2019). EVs obtained from multipotent mesenchymal stem cells (MSC) are of particular interest for therapeutic use (Lener et al., 2015) since they convey anti-inflammatory (Romanelli et al., 2019) and anti-apoptotic effects (Terlecki-Zaniewicz et al., 2018), stimulate angiogenesis (Zhang et al., 2019) and wound healing (Shabbir et al., 2015). A current clinical study investigates the use of MSC-EVs for the treatment of type I diabetes mellitus (NCT02138331). In an individual treatment attempt, MSC-EVs were administered in a patient with graft versus host diseases (Kordelas et al., 2014). The properties of an EV-based therapeutic, a product of natural processes, fluctuate widely due to the variable cellular secretome. Furthermore, the amount and composition of EVs that can be produced is impacted by the cell cultivation and expansion conditions as well as by the EV harvesting and isolation techniques, e.g. the media supplements used during MSC cultivation (Gimona et al., 2017; Konoshenko et al., 2018). From the cellular secretome, a mixed population of EVs, such as exosomes, microvesicles, shedding vesicles (ectosomes), and microparticles, as well as a number of soluble factors, all of which are also involved in intercellular communication (Gurunathan et al., 2019), can be enriched. Reports indicate that also shedding vesicles, released from the cell membrane, participate in important physiological and pathological processes such as coagulation, inflammatory diseases, and tumor progression (Cocucci et al., 2009). The definition and control of EV-based therapeutics is subject to the guidelines of the regulatory network for medical products. Although EVs were used in patients for the first time in 2005 for cancer therapy (Escudier et al., 2005), there are still no routinely established standard techniques for quality control of EV therapeutics for clinical production. The latest minimal information for studies of extracellular vesicles (MISEV) guidelines stated that quantitation and single-particle characterization should be performed by methods including but not limited to sizing and counting by particle tracking techniques, imaging by electron microscopy, and advanced flow cytometry (Witwer et al., 2019). Current particle quantitation methods such as nanoparticle tracking analysis (NTA), dynamic light scattering (DLS), and resistive pulse sensing (RPS) cannot discriminate EVs from other particles (Giebel and Helmbrecht, 2017). To detect EV surface proteins, novel techniques are available, including laser tweezers Raman spectroscopy (Carney et al., 2017) or surface plasmon resonance (SPR) (Rupert et al., 2016). While these techniques provide information such as the number of particles or the presence of characteristic surface proteins, they cannot distinguish between EVs and other particles (e.g. protein aggregates) which may be enriched during isolation (Lobb et al., 2015), so analysis results remain ambiguous. Due to their heterogeneity and small dimensions, EV characterization is still challenging. AFM imaging can be used to identify the influence of EV isolation regarding size distribution, morphology and mechanical properties (Parisse et al., 2017). Perissinotto et al. analyzed EVs with a combination of Fourier Transform Infrared Spectroscopy, Ultraviolet Resonant Raman Spectroscopy, AFM, and Small Angle X-Ray Scattering to address size, stability and purity of EVs (Perissinotto, 2020). A more advanced method for measuring particle number combines SPR with atomic force microscopy (AFM; Obeid et al. (2017)). The latter presents a promising first strategy to obtain more detailed information about EV properties (Vorselen et al., 2020). Nevertheless, ensuring that the counted particles are indeed EVs and that these EVs are therapeutically active (e.g. as indicated by their surface marker profile and meaningful biochemical or cell-based assays), in a robust, accessible and economical manner remains unsolved. Quartz crystal microbalance (QCM) is a highly sensitive method able to simultaneously detect biomarker association and the resulting changes in mechanical properties at the binding interface (Suthar et al., 2020). QCM provides highly sensitive mass sensing, based on the piezoelectric effect via changes in the oscillation of a gold-coated quartz crystal. It is a recognized and established method for the detection and characterization of adsorption processes on surfaces. Changes in frequency directly relate to changes in mass according to the Sauerbrey equation (Sauerbrey, 1959), given thin and rigid adlayers. Soft and thus more dissipative adlayers such as those represented by biological binding processes often require more complex modeling (Reviakine et al., 2011). Specific interactions of ligand-receptor or antigen-antibody as well as the behavior of supramolecular systems such as the adhesion of vesicles and liposomes can be examined with a sensitivity of less than 1 ng/cm^2^ in real time to determine kinetic rates and affinity constants (Kasper et al., 2016; Strasser et al., 2020). The simultaneous detection of energy loss during the oscillation of the quartz crystal – adsorbate system allows experimental access to interfacial changes that do not alter the mass of the system but do affect its rigidity. It enables the differentiation between rigid and soft adsorbates, and - using the state of the art QCM-D device – has been employed previously to differentiate between vesicle deposition to functionalized sensors and artifacts (Suthar et al., 2020). Here we propose a QCM-I and AFM based protocol to characterize EVs tailored to be robust, reproducible, and easy to integrate in existing laboratory infrastructures.

## Methods

### Preparation and characterization of hUC-MSC and MSC-EVs

#### Primary isolation and expansion of human mesenchymal stromal cells

Human umbilical cord (UC)-derived MSC were isolated as previously described (Pachler, 2017). Immediately after delivery, cords were collected and stored in phosphate buffered saline (PBS) until further processing. Whole cords were washed with PBS to remove contaminating blood cells before the cord stroma was cut into small pieces of 1 – 2 mm^3^. Pieces were transferred into a culture plate allowing them to dry-adhere to the plastic surface before adding culture medium based on alpha-modified minimum essential medium (α-MEM Sigma-Aldrich, MO, USA) supplemented with 10% (v/v) pooled human platelet lysate (pHPL) and Dipeptiven (5.5 mg/ml, Fresenius-Kabi, Germany). Pooled HPL was prepared as previously described (Laner-Plamberger et al., 2015). In brief, expired irradiated platelet concentrates were lysed by several freeze/thaw cycles. Platelet fragments were pelleted by centrifugation (4000 x *g,* 15 min at room temperature) and aliquots of the supernatant were frozen at −30°C until use. After 10 to 12 days, outgrowing UC-MSC colonies became visible and cord tissue pieces were removed. UC-derived MSC were detached enzymatically by addition of TrypLE Select CTS (A12859-01, Thermo Fisher Scientific, MA, USA), and further expanded in cell factory systems (CF4, Thermo Scientific). Immuno-phenotyping and viability analysis of MSC was carried out according to the proposed surface marker profile for defining MSC identity as published by the International Society of Cell Therapy (ISCT) in 2006 (Dominici et al., 2006).

#### Manufacturing and characterization of MSC-EVs

Batches of EVs from UC-MSC were prepared according to Good Manufacturing Practice (GMP) as previously described (Desgeorges et al., 2020; Gimona et al., 2017; Pachler et al., 2017b). In brief, cells were cultured in fibrinogen-depleted culture medium at 5 % CO_2_ and 37°C. Upon reaching 60-70% confluence, growth medium was exchanged with EV-depleted harvest medium containing 5% HPL. After 24 h, conditioned medium was collected, centrifuged and purified via sterile filtration (0.22 μm). The resulting supernatant was concentrated and buffer-exchanged to PBS by Tangential Flow Filtration (TFF) and diafiltration, respectively, using a 100 kDa hollow fibre column filter (Spectrum Labs, Greece). Ultimately, EVs were enriched by ultracentrifugation at 120.000 x g for 3 h at 18 °C in a Sorvall model WX-80 using a fixed angle rotor model Fiberlite F37L-8×100 and the resulting pellets were resuspended in Ringer’s Lactate and again sterile filtered. Individual doses were stored in glass vials at – 80 °C and batches were tested for endotoxin levels, bacterial sterility and the presence of mycoplasma.

#### Total protein mass determination

Total protein amounts were determined using a QuBit® 3.0 Fluorometer instrument (Life Technologies, CA, USA) according to the manufacturer’s instructions.

#### Cytokine profiling

Cytokines (IFN-gamma, IL-10, IL-12p70, IL-13, IL-1β, IL-2, IL-4, IL-6, IL-8, TNFod, β-NGF and BDNF) from various preparations were analyzed using V-Plex and U-Plex human multiplex immunoassay kits on the MSD platform (Meso Scale Diagnostics, MD, USA) according to the manufacturer’s instructions.

#### Nanoparticle tracking analysis (NTA) in light scatter mode

To determine the size and amount of particles in the individual EV preparations, samples were analyzed using a Nanoparticle Tracking Device (ZetaView PMC 110 from Particle Metrix, Germany) in light scatter mode essentially as described (Desgeorges, 2020). Previously frozen EV preparations were used and samples were diluted to a concentration of 4 – 7 x 10^7^ particles/mL in PBS. Prior to NTA analysis, the instrument was calibrated using Yellow/Green-labeled 100 nm polystyrene standard beads (1:1.000.000 dilution in ddH_2_O). The minimum brightness was set to 20 arbitrary units (AU), temperature to 21.5 °C, shutter to 70 AU, and sensitivity to 85 AU. Subsequently, data for two exposures at 11 measurement positions were acquired per sample. Based on the Stokes-Einstein equation, particle size was calculated using the ZetaView software (PMX 110: Version 8.4.2).

#### MACSPlex surface protein profiling

The bead-based multiplexed FACS-based MACSPlex Exosome Kit (Miltenyi Biotec, Germany) is an assay for the analysis of surface markers present on EVs. To characterize the various MSC-EV preparations we used the MACSPlex kit according to the manufacturer’s instructions and following a validated standard operating procedure with 5 x 10^7^ to 5 x 10^8^ total particles as input. Data acquisition was done using a FACS Canto II instrument (BD Biosciences, CA, USA). For additional CD73 analyses an anti-CD73-BV421 antibody (BD Biosciences) was used. Data normalization was directed towards CD9/CD63/CD81 APC signal. Isotype control normalization was performed as described (Wiklander et al., 2018).

#### CryoEM analysis

UC-MSC-EV samples were deposited on an electron microscopy (EM) grid coated with a perforated carbon film. Samples were quickly frozen in liquid nitrogen-cooled liquid ethane using a Leica EM-PC cryo system. EM grids were maintained under liquid nitrogen until use. EM grids transferred to a Tecnai F20 cryo-electron microscope (FEI, ThermoFisher) operating at 200 kV. Grids were mounted in a Gatan 626 cryo-holder and images were recorded with a FEI-Eagle camera.

#### CD73 activity assay

The enzymatic activity of CD73 in UC-MSC-EV preparations was determined by incubating 10 μL of EVs in 10 mM HEPES (Sigma H3537) buffer containing 2 mM MgCl_2_ (Merck Millipore, MA, USA) with 10 μM AMP (Sigma 01930) for 20 min at 37°C. The amount of AMP consumption was detected with the AMP-Glo™ Assay Kit (Promega, WI, USA) according to the manufacturer’s protocol and measured with Spark^®^ multimode microplate reader (Tecan, Austria). 2 ng rhCD73 (Sigma N1665) was used as positive control and AMP-CP (Sigma M8386) as CD73 inhibitor.

#### Immunomodulation assay

To investigate the immunomodulatory activity of UC-MSC-EV preparations, the capacity to inhibit T cell proliferation in vitro was studied, essentially as described previously (Pachler et al., 2017a). Stimulation of T-cell proliferation was achieved by incubation with CD3/CD28 antibody beads (Thermo Fisher). Carboxyfluorescein succinimidyl ester (CFSE) pre-labeled pooled peripheral blood mononuclear cells were stimulated with CD3 and CD28 and co-cultured with different ratios of UC-MSC-EVs for 72 hours essentially as described (Trickett and Kwan, 2003). The percentage of inhibition of fluorescently labeled CD3 T-cell proliferation was analyzed by flow cytometry in triplicates.

### QCM

Quartz crystal microbalance (QCM) is an ensemble technique utilizing the inverse piezoelectric effect to measure mass adsorption and changes in fluid viscosity. In QCM, adsorption processes and changes in the medium surrounding the crystal result in in proportional changes in the frequency and energy dissipation of its oscillation (Cheng et al., 2012). Monitoring changes in resonance frequency due to mass deposition on the quartz surface thus yields real-time traces of association and dissociation of everything from small molecules to entire cells.

#### Lipid preparation

Liposomes containing biotinylated headgroups were prepared from 1,2-dioleoyl-sn-glycero-3-phosphoethanolamine-N-(cap biotinyl) (Biotinyl-cap-DOPE), 1,2-dioleoyl-sn-glycero-3-phosphocholine (DOPC) and 1,2-dioleoyl-sn-glycero-3-phospho-L-serine (DOPS) in a ratio of 1:7:2 (Karner et al., 2017). In total 5 mg of lipids (all Avanti Polar Lipids, AL, USA) of Biotinyl-cap-DOPE:DOPC:DOPS were dissolved in 2 mL of Trichlormethan/Chloroform (Roth, Germany) and mixed with 1 mL of Methanol (Roth, Germany). Lipids were dried using a R-100 rotary evaporator (Buchi, Germany) at 180 rpm for 30 min. To remove the methanol, the dried lipid mixture was dissolved in 3 ml chloroform and dried again using the rotary evaporator at 180 rpm for 30 min. Afterwards, the lipid mixture was further dried for 1-2 hours with a high vacuum pump DCP 3000 (Vacuubrand, Germany). The dried lipid mixture was dissolved in 500 μL mQ-H_2_O by repeated pipetting and transferred into a glass vial capped with a septum. The lipid mixture was sonicated in an ultrasonic bath (Sonorex, Germany) for 10 min until the emulsion got clear. To obtain a final concentration of 2 mg/ml, 2 mL of running buffer (RB) consisting of 10 mM Hepes, 150 mM NaCl, 2 mM CaCl_2_ (pH 7.4) were added and aliquots of 100 μL were shock-frozen with liquid nitrogen and stored at −80°C until further use.

#### QCM measurements

All QCM experiments were conducted using a two-channel QCM-I system from MicroVacuum Ltd. (Hungary). AT-cut SiO_2_-coated quartz crystals with a diameter of 14.0 mm and a resonance frequency of 5.000 MHz were used (Quartz Pro AB, Sweden). All sensorgrams were recorded on the first, third and fifth harmonic frequencies simultaneously. The data shown relates to the third harmonic. A constant flow rate was provided by a programmable peristaltic pump. Before each set of experiments, the SiO_2_-coated QCM crystals were cleaned as described (Strasser et al., 2020). Briefly, chips were immersed in 2% SDS for 30 min, followed by thorough rinsing with mQ-H_2_O. The chips were then dried in a gentle stream of N2 and activated using air plasma (4 min at 80 W), after which they were ready to be mounted in the measurement chamber. Once securely affixed, the sensor surface was cleaned by pumping 2% SDS through the liquid cell at 250 μL/min for 5 min, followed by mQ-H_2_O at 250 μL/min for 5 min, and finally RB for equilibration at 50 μL/min. All subsequent injections were performed at 50 μL/min. After approximately 30 min of RB equilibration, 200 μg/ml biotin-liposome solution was injected until complete bilayer fusion was achieved (approx. 20 min). To ensure complete bilayer coverage without exposed substrate, a 6.25 μg/ml Control Protein (CP) solution was injected. The used CP, C1q (Complement Technology, TX, USA), adheres to activated SiO_2_ but not to the lipid bilayer. No change in frequency proves that the lipid is closed and that there is no non-specific protein adsorption. Subsequently the bilayer is loaded with 60 μg/ml streptavidin (Merck Millipore, MA, USA) to saturation. Finally, 10 μg/ml biotinylated antibody are added and bound to the lipid bilayer via streptavidin. Specific EV markers were recruited using biotin anti-human IgG1 targeting CD9 (312112), CD44 (103003), CD63 (353017), CD73 (344017) and CD81 (349514) (all from BioLegend, CA, US), CD49e (Invitrogen 13-0496-80) respectively. Biotin anti-mouse IgG1 isotype control (400104) served as negative control (BioLegend, USA). Then UC-MSC EVs can be analyzed for the presence of different surface markers by diluting the stock concentration 1:10 with RB and injecting the solution for 5 min to ensure appropriate volume exchange. After this time the flow was stopped for 30 min in order to limit sample consumption to <300 μL and to allow for stable EV association to the surface. Finally, the sample was washed with RB for 20 min under constant flow to observe any dissociation of the loaded EVs. Finally, the EV recruitment efficiency, i.e. the frequency shift during EV binding normalized to the shift during antibody immobilization, is calculated and compared between markers using MATLAB (MathWorks). Analysis of frequency and dissipation response was performed with the QCM software BioSense, version 3.11.

### AFM

An atomic force microscope (AFM, JPK Nano Wizard4, Germany) mounted on an Olympus IX71 inverted optical microscope was used for sample characterization. MLCT-F cantilevers with a nominal tip radius of 40 nm and a spring constant of 0.6 N/m (Bruker, Germany) were used for measurements. Cantilevers were calibrated prior to each measurement using the commercial JPK NanoWizard control software V6.1.163. The indentation force was set constant to a value of 0.8 nN for EV-imaging.

#### Imaging Mode

The off-resonance JPK QI™ mode was used for topography imaging and mechanical characterization of EVs. In this mode, several sample properties (topography, stiffness, adhesion, elasticity) can be acquired within a single measurement by recording complete force-distance curves in each pixel. As a result, lateral forces on the sample are eliminated during scanning which minimizes sample deformation and possible displacements for loosely bound substances. All measurements were done in an aqueous environment (PBS).

#### Sample preparation

Glass slides were cleaned by rinsing with mQ-H_2_O, Isopropanol (99,9%), mQ-H_2_O and Ethanol (96%), and dried with a stream of N2 gas. Glass slides were cleaned in Argon plasma for 15 min at 40 kHz 96 W (Zepto, Diener electronic Germany). Samples were diluted in PBS, until a sparse surface density of ~2 particles/μm^2^ was achieved after incubation for 45 min on the cleaned glass slide to ensure proper separation between objects for individual characterization. The sample were washed 3 times with PBS and measured on the same day.

#### Quantitative Analysis

For differentiation between the species detected on the surface, topography and elasticity images have been used. Particles with a height of >8 nm above the background that present with a clearly delineated lower elasticity than the substrate have been classified as EVs, as previously described using tapping mode phase imaging (Sharma et al., 2010) and force-distance curve-based deformation imaging (Vorselen et al., 2020). Other particles were designated as non-vesicular particles (NVPs). For characterization, the height and aspect ratio (height/width at FWHM) were extracted from AFM images using JPK data processing software, version 6.1.121.

## Results

### Quality control of UC-MSC-EVs according to MISEV guidelines

In this study, UC-MSC-EVs that were manufactured according to the pharmaceutical standards and guided by approved Standard Operating Procedures (SOPs) were enriched from the MSC secretome by Tangential Flow Filtration (TFF) and ultracentrifugation (UC). The MISEV 2018 guidelines state that both, the source and the preparation of EVs, should be described as quantitatively as possible. First, the number of cultured cells at the time of collection must be determined for each experimental use (Thery et al., 2018). As shown in Table 1, isolated MSC exhibit a viability of ≥ 90% and represent the typical MSC surface marker profile with ≥ 95% positive for CD 29+, CD44+, CD73+, CD90+, CD105+, CD166+ and with ≤ negative for CD14-, CD19-, CD34-, CD45- and MHC class II-. Regarding quantitation and single-particle characterization of EVs, particle tracking, imaging, and advanced flow cytometry has been recommended (Witwer et al., 2019). Our NTA resulted in a particle number of > 5×10^10^/ml, particle sizes ranging from 80 to 150 nm (Table 1, Fig. 1a), and a fraction ≥10-15% of CD63+, CD81+ and CD73+ EVs. Additionally, a total protein concentration of <5 mg/ml was determined (Table 1). Analyzing these EVs using MACS Plex, we observed a panel of positive (CD9+, CD29+, CD44+, CD49e+, CD63+, CD81+, CD73+, CD105+, MCSP+) and negative (CD14-, CD19-, CD34-, CD45-, CD142-, CDMHC class I-, class II-) surface markers that are typical for MSC-EVs (Table 1, Fig. 1b). The cytokine profile of MSC-EV preparations demonstrates the absence of pro-inflammatory cytokines IFN-gamma, IL-10, IL-12p70, IL-13, IL-1β, IL-2, IL-4, IL-6, IL-8, TNF-α, β-NGF and BDNF (Fig. 1c). Further characterization of the MSC-EVs was performed using cryo-electron microscopy, where the lipid bilayer of EVs was confirmed (Fig. 1d), and AFM, where EV height was correlated with their size, an indication of the homogeneity of the sample (Fig. 1e).

**Figure 1:**
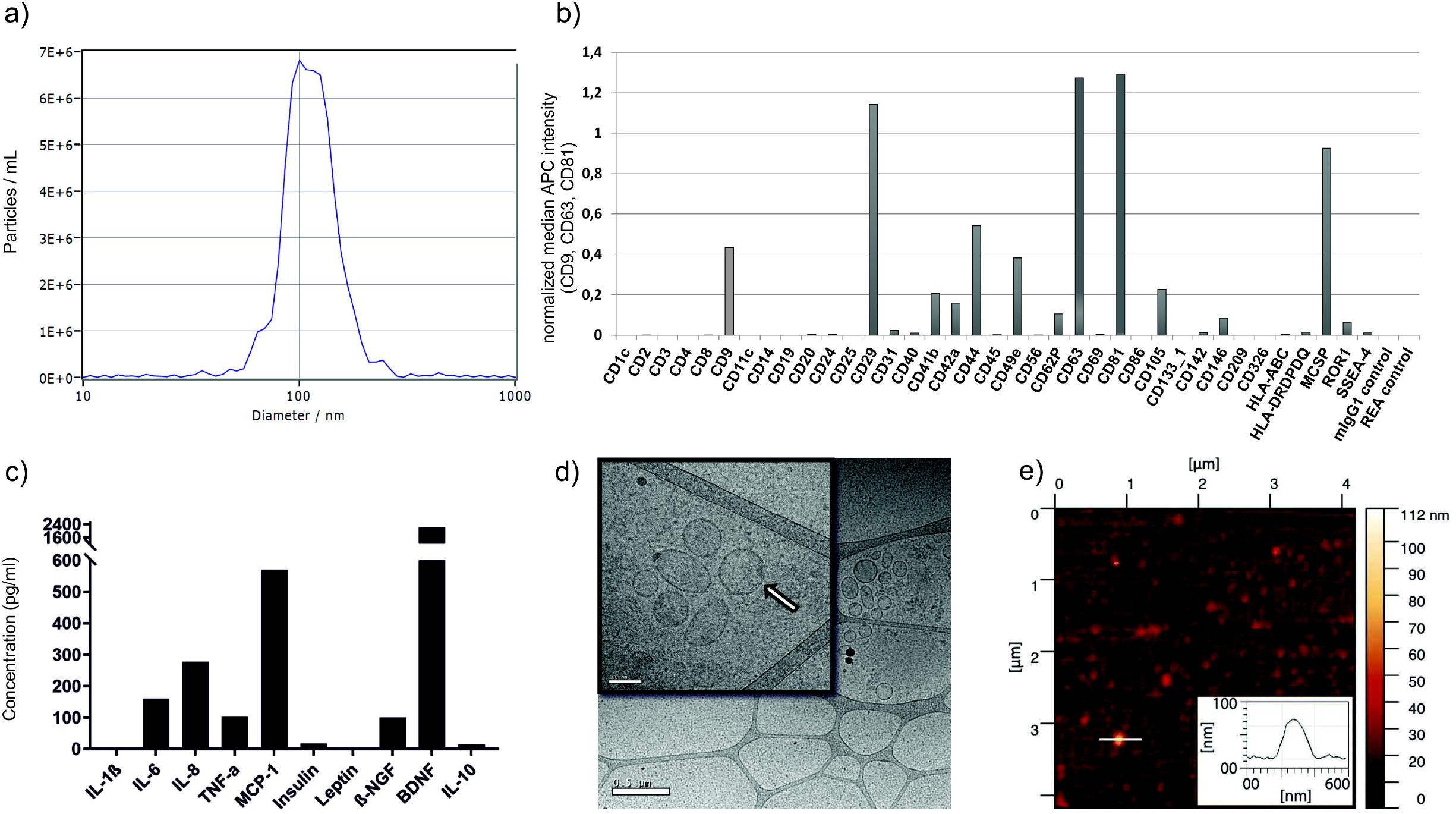
Characterization of UC-MSC-EVs. (a) Size distribution of UC-MSC-EVs by nanoparticle tracking analysis. (b) Surface marker profiling by MACSPlex. (c) UC-MSC-EV cytokine profile revealed absence of pro-inflammatory cytokines by multiplex analysis. (d) Cryo-electron microscopy image from a representative batch of MSC-derived EVs, size bar 0.5μm. The lipid bilayer surrounding the EV can be recognized (arrow in the inset; size bar 100nm). (e) AFM topographic image and one corresponding line profile (inset) of UC-MSC-EVs.

**Table 1:**
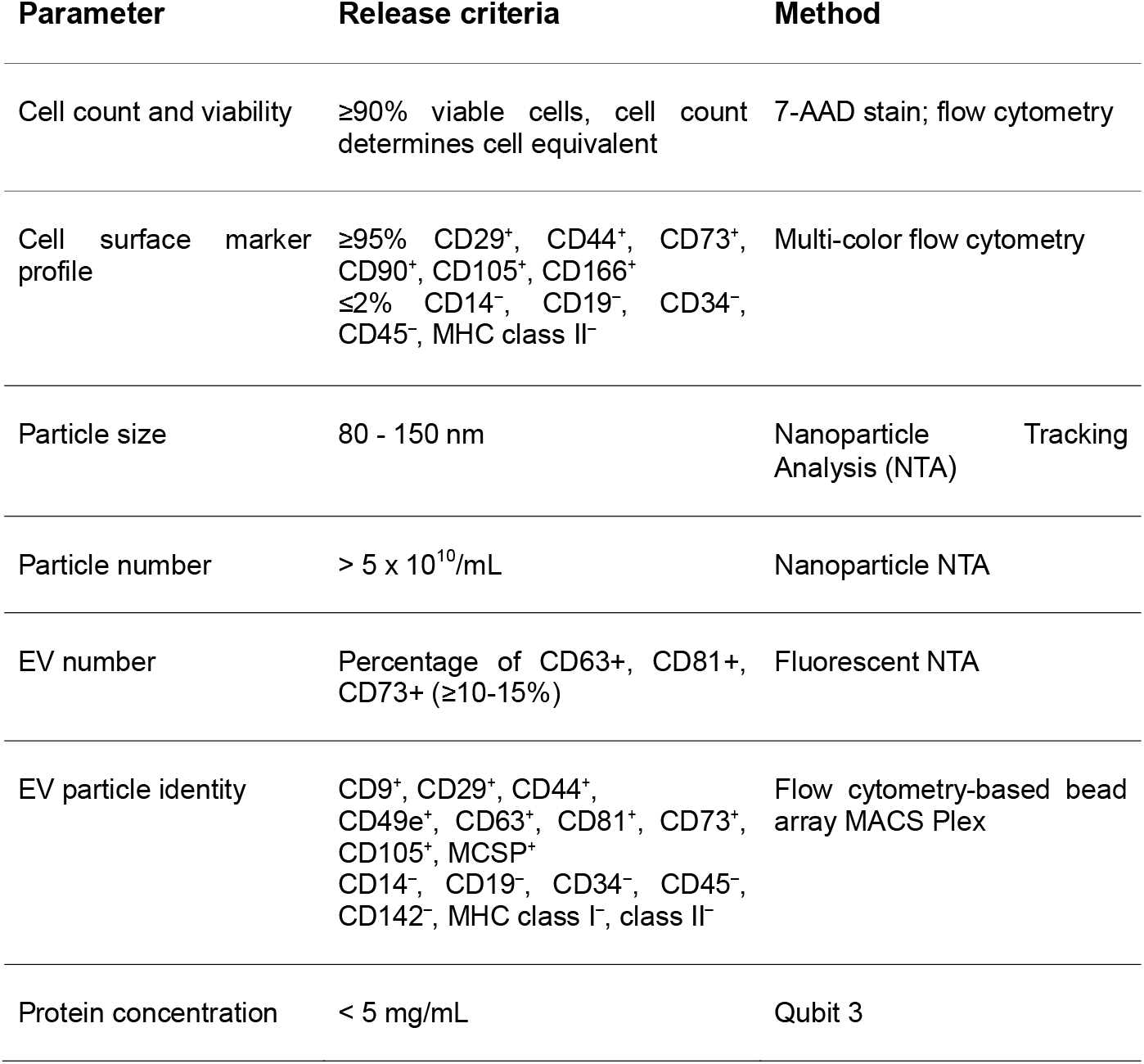
Multimodal Quality Control Parameters of UC-MSC EVs. Parental cell characterization, identity, purity and impurity determination of EV preparations is performed for the standard quality release testing of all research scale preparations and for GMP training and GMP clinical runs.

### Functional characterization of UC-MSC-EVs

We have shown previously, that UC-MSC-EVs retain the immunomodulatory potential of their parent MSCs (Pachler et al., 2017a). The precise mechanism underlying the T-cell proliferation inhibition and potential immunomodulatory activity of UC-MSC-EVs is not completely understood. However, adenosine signaling via the conversion of AMP to adenosine by the GPI-anchored 5’-ecto-nucleotidase CD73 can shape various lymphocyte functions, the majority of effects being described as suppressive, and could be, at least in part, responsible for this activity. We thus investigated if the EV-associated CD73 is still functional on enriched UC-MSC-EVs after the purification process. As shown in Figure 2 UC-MSC-EV preparations actively converted AMP in a biochemical assay in a dose dependent manner. This activity was sensitive to heat and heat-treated UC-MSC-EV preparations which were neither capable of converting AMP nor of inhibiting T-cell proliferation in vitro. These findings suggest that the UC-MSC-EVs that were used in this study retained biochemical and biological activity.

**Figure 2:**
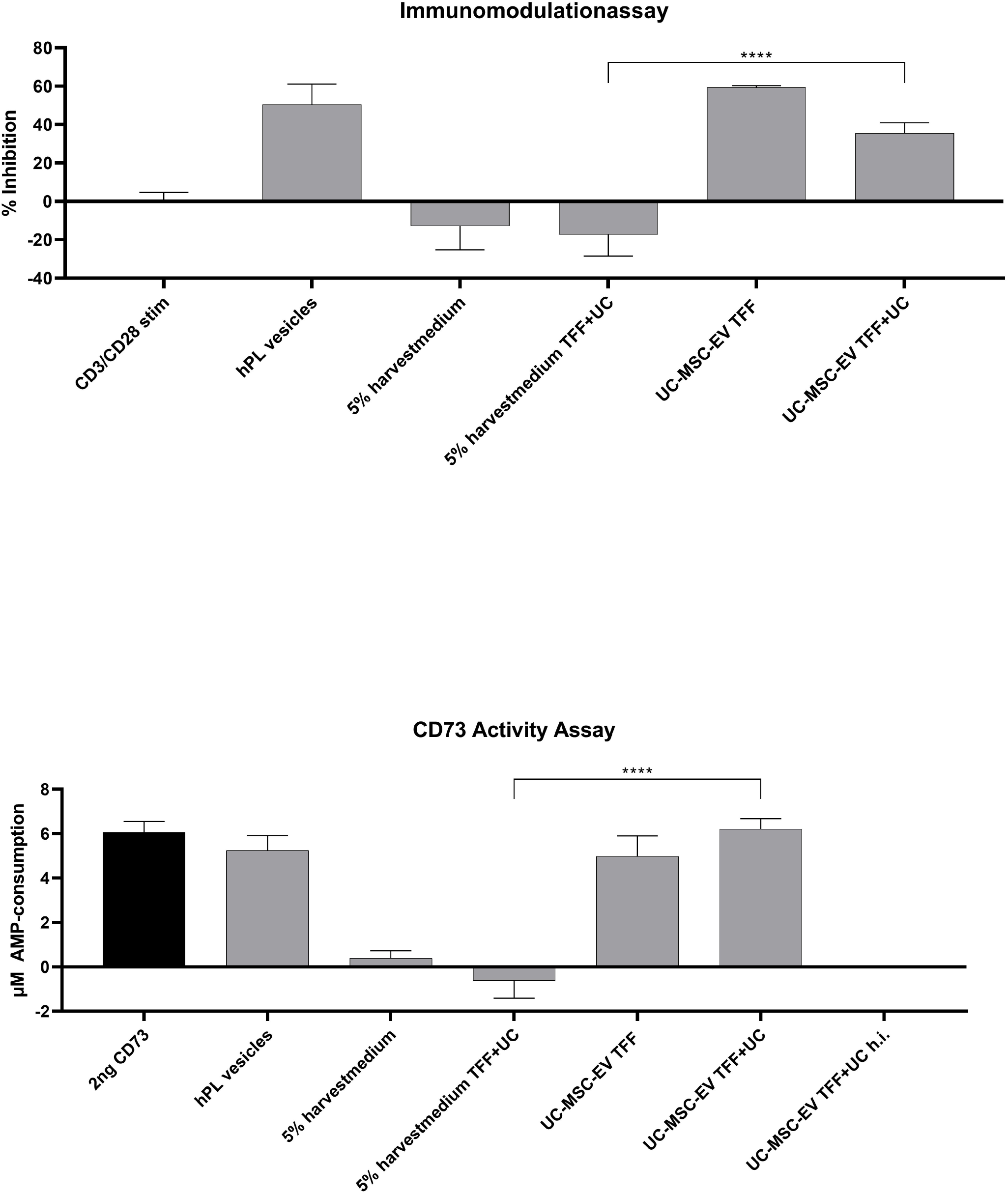
Characterization of biochemical and biological activity of CD73-positive UC-MSC-EVs. (a) Immunomodulation. CD3/CD28-stimulated pre-labeled pooled human peripheral blood mononuclear cells were incubated with UC-MSC-EV preparations at different steps of the manufacturing process (TFF alone (TFF) or TFF followed by ultracentrifugation (TFF + UC)). The T-cell proliferation-inhibiting activity residing in the growth supplement human platelet vesicles (hPL vesicles) was abolished in the vesicle-depleted harvest medium. Subjecting the harvest medium to the same purification procedure as used for the MSC-derived EVs (5% harvest medium TFF + UC) did not accumulate or restore any inhibitory activity. (b) EV-associated CD73 is biochemically active. The AMP-converting activity that is present in the growth supplement human platelet vesicles (hPL vesicles) was abolished by vesicle depletion of the harvest medium by tangential flow filtration. Subjecting the harvest medium to the same purification procedure as used for the MSC-derived EVs (5% harvest medium TFF + UC) did not accumulate or restore the enzymatic activity. UC-MSC-EV preparations containing CD73-positive EVs exhibit significant enzymatic activity after both TFF and TFF+UC purification. By contrast, heat inactivation of UC-MSC-EVs for 5 minutes at 95°C abrogated the enzymatic activity of CD73 (UC-MSC-EVs TFF + UC-h.i.). As a control 2 ng recombinant human CD73 was used.

### Advanced AFM characterization

AFM is a well-established technique for high-resolution imaging of biological structures under physiological conditions (Preiner et al., 2015; Preiner et al., 2014; Preiner et al., 2007; Strasser et al., 2019). Beyond simple EV height profiles, AFM can be used to determine the mechanical properties of EVs (LeClaire, 2020). Herein, we used AFM imaging to map the elasticity of UC-MSC-EVs adsorbed to glass slides and correlated this property with particle sizes (aspect ratios). Together, these data allowed us to perform a quantitative analysis of the samples at single particle level and to distinguish EVs from other NVPs (Fig. 3a and b). Due to their overall similarity in morphological appearance, distinction between EVs and NVPs based on topography images alone was not possible. Recent studies reporting on AFM phase-imaging of EVs on rigid supports suggest that the vesicles present with a rigid outer rim and a soft central region, resulting in “doughnut-like” tapping mode phase profiles (Sharma et al., 2010). Accordingly, simultaneous analysis of topography and QI™-mode elasticity maps (Fig. 3a, b) revealed two distinct populations of particles that differ in their specific elasticity profiles, i.e. becoming increasingly softer from the periphery to their center (Fig. 3b, profile 2, Supplementary Fig. 1), or exhibiting a more homogenous elasticity hardly distinguishable from the background (Fig 3b, profile 1). This enabled us to identify the former population as EVs, while the latter population lacking these specific properties are consequently classified as NVPs. In this way, we analyzed 84 nanoparticles and identified 41 EVs and 43 NVPs (Supplemental Fig. 1).

**Figure 3.**
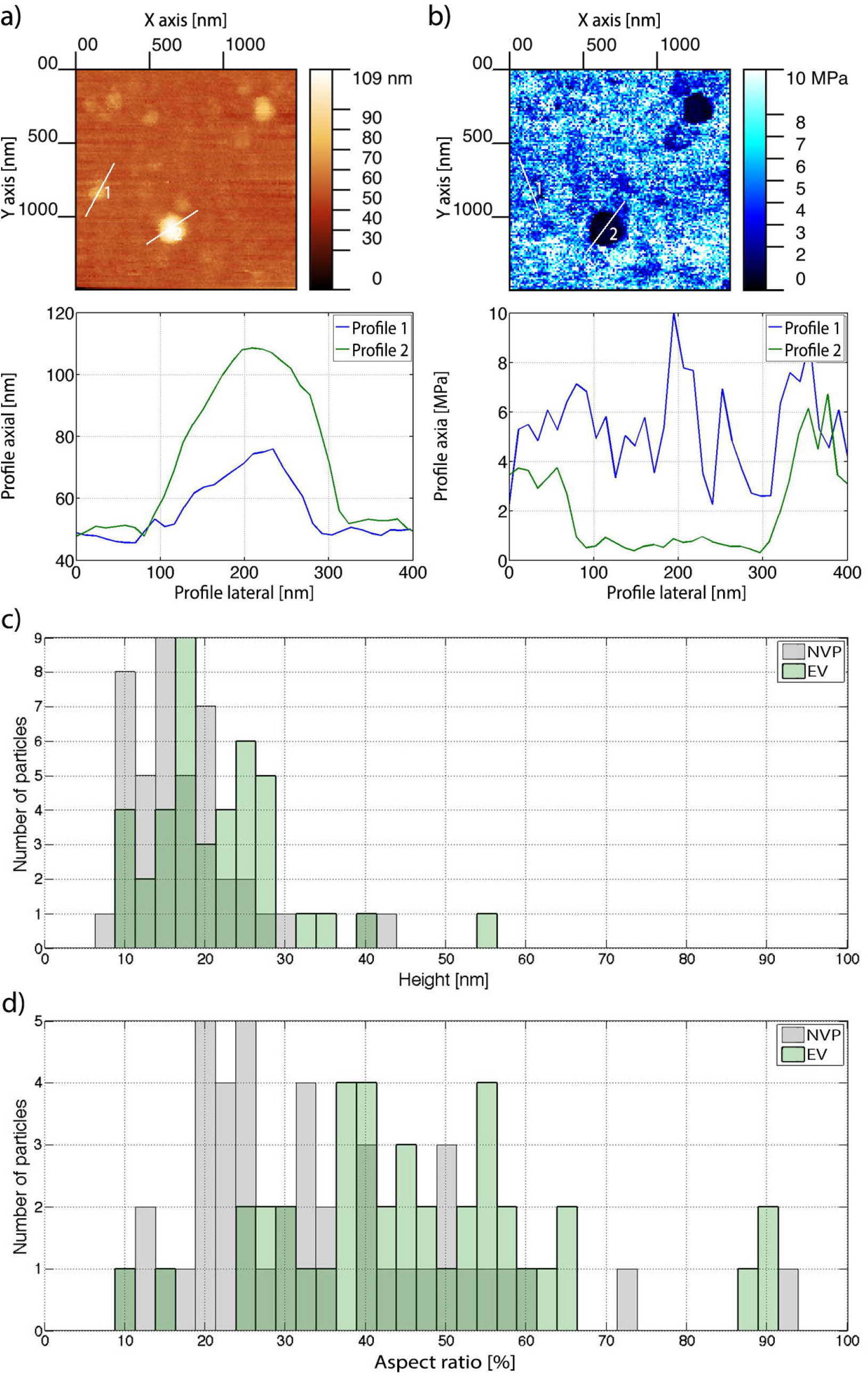
Exemplary AFM image of EVs. (a) Topography image, showing particles of various sizes (top). The corresponding height-profiles of the two marked particles are displayed below, (b) Simultaneously recorded elasticity map of the same sample area as in (a) (top). EVs are distinguished from non-vesicular particles (NVP) by clearly delineated drop in Young’s Modulus (profile 2) whereas other particles exhibit a constant elasticity indistinguishable form the background (profile 1). (c, d) Histograms of particle size distributions and the lateral to axial size aspect ratios determined via AFM. (c) Height distribution of NVPs (gray) and EVs (green). (d) Histogram representing aspect ratios (height/width) of NVPs (gray) and the EVs (green). N=84.

Fig. 3c and d depict the distribution of particle heights and aspect ratios (ratio of FWHM in scanning direction and height) for these two populations with mean heights of 21.5 nm ± 8.8 nm and 17.3 nm ± 7.3 nm as well as average aspect ratios of 46 ± 18 and 34 ± 17 for EVs and NVPs respectively.

To test whether these empirical distributions where drawn from the same probability distribution, we performed a parameter-free Kolmogorow-Smirnow-Test (KS-Test) which resulted in highly significant asymptotic p-values of 2*10^-2^ for height (Fig. 3c) and 10^-3^ for aspect ratio (Fig. 3d). Thus, the geometrical properties of EVs and NVPs can be considered distinct, which potentially offers a way to determine the percentage of EVs included in a similar EV preparation solely from topographical data by fitting a linear combination of the respective distributions for EVs and NVPs as determined in our QI™-mode experiments to the aspect ratio distribution of all particles (highest significance) of the unknown sample. However, reliable distinction of individual objects solely based on topography is impossible, making elasticity mapping via QI™-mode imaging or AFM tapping-mode phase images an essential tool to characterize heterogeneous EV-NVP formulations at the single-particle level.

### QCM measurements

Our results so far describe a biologically active and potentially therapeutically relevant but complex mixture of EVs and NVPs. We thus established a flexible and easy to use QCM-I protocol for the detection of surface markers on the EV fraction, which utilizes a combination of specific EV recruitment by antibodies and the high viscoelasticity of fluid-filled vesicles as compared to stiff protein aggregates. Figure 4a provides a schematic illustration of the experimental setup. At first, a lipid bilayer containing 10% biotinylated lipids was established on the QCM sensor chip (Fig. 4b). The corresponding characteristic total frequency shift amounted Δf_Lipid_ of ~25-30 Hz (Keller and Kasemo, 1998). We employed a 450 kD control protein, which adsorbs to exposed SiO_2_ but not to lipids (Fig. 4c) to check for bilayer integrity and prevent unspecific binding of antibodies and EVs later in the experiment. The surface was then loaded with streptavidin and subsequently with the biotinylated IgG of interest (Δf_SA_ and Δ_IgG_ respectively, both ~20Hz). Finally the EV solution was injected, resulting in a maximal association signal Δf_EV,max_ and, after a dissociation phase, a stable immobilization signal Δf_EV stable_. CD44 exhibited the largest recruitment efficiency (i.e. the frequency shift during EV binding normalized to the amount of immobilized IgGs on the surface, Δf_EV,max_/Δf_IgG_) followed by CD49e, CD63, CD81, CD73 and CD9, while the negative control resulted in negligible unspecific recruitment (Fig. 5a). Thus, antibodies targeting common EV markers, the tetraspanins CD9, CD63 and CD81, showed significant recruitment of the UC-MSC-EV sample relative to the control, as did the typical MSC markers CD44 and CD73, and the marker typical for UC-MSC, CD49e (Fig. 5a). This is in line with our EV surface marker expression characterized by MACSplex analysis (Table 1, Fig. 1). Dissipation monitoring revealed an increase in energy dissipation directly after EV injection for all markers except the negative control. The observed increase (in the range of 2-9×10^-6^) marks the bound mass as highly viscoelastic (Cho et al., 2007) and thus with high probability as EVs, rather than rigid particles or protein aggregates (Fig. 5b). This approach to characterize EV surface markers additionally provides a semi-quantitative comparison of EV recruitment: Markers that result in recruitment efficiencies below that of the negative control plus 3 standard deviations (n.c.+3) are considered negative, those with values between n.c.+3 and n.c.+5 as weakly positive, and those above n.c.+5 as strongly positive (Fig. 5c). Given these parameters, CD9 is classified as weakly positive, while CD81, CD73, CD63, CD49e, and CD44 are strong positives. No major distinction can be made between relative EV recruitment efficiencies with or without 20 min of dissociation (Fig. 5d). An additional parameter provided by all QCM devices that simultaneously record frequency and dissipation is the change of these two parameters relative to each other, ΔD/Δf, providing a straightforward way to compare the viscoelasticity of adsorbates (Fig. 5b inset; (Saftics et al., 2018; Tagaya, 2015)). The high ΔD/Δf ratios for all investigated vesicle samples (fluid phase liposomes, gel phase liposomes, MSC-EV formulation) adsorbing to bare SiO_2_-coated chips indicates their viscoelastic nature. In comparison, the control protein results in significantly lower changes in dissipation, marking it as more rigid. Direct comparison of dissipative losses on SiO_2_ showed highest elasticity with fluid-phase liposomes (DOPC), followed by gel phase liposomes (DPPC) and our UC-MSC-EVs where a moderate viscoelasticity could be observed. This fact, coupled with the specificity of recruitment due to our antibody-based EV immobilization approach, provides a high degree of confidence that EVs, not NVPs, were measured. With this QCM setup, we can thus capture specific EVs with a respective therapeutic functional marker.

**Figure 4.**
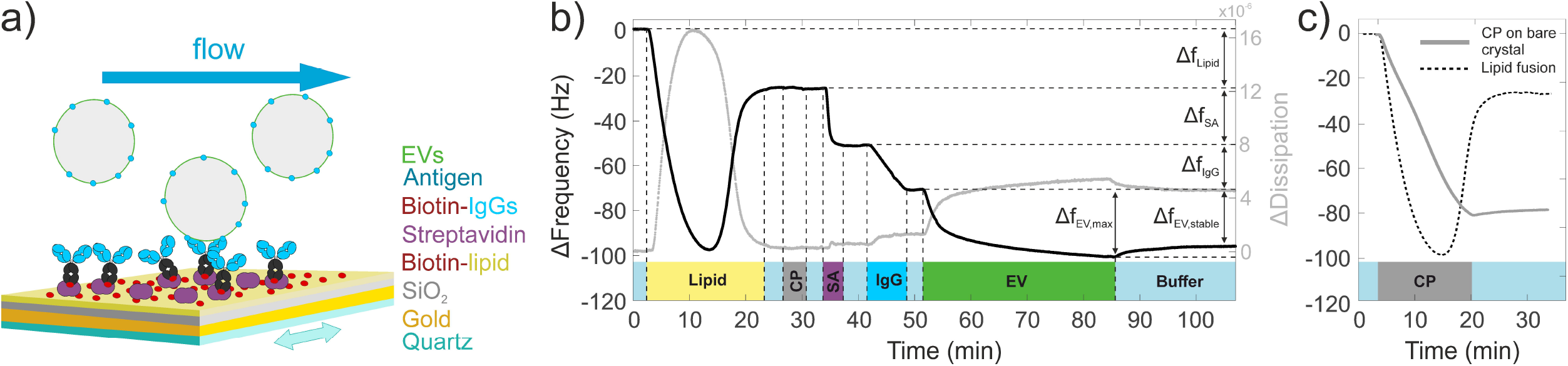
QCM workflow for characterizing EV surface proteins. (a) Schematic of the experimental setup. A quartz crystal is coated with a supported lipid bilayer containing biotin. Streptavidin is added and recruits biotinylated antibodies, which subsequently capture antigen-carrying EVs. Vesicles, extracellular particles, or proteins without this marker are not detected. (b) Representative QCM sensorgram of an EV characterization experiment. Δf_Lipid_ serves as indication of lipid coverage and quality, as does the subsequent injection of the Control Protein CP. No binding of CP was observed after lipid fusion, demonstrating that a homogenous lipid bilayer without defects or non-specific protein adsorption has formed. Δf_SA_ and Δ_IgG_ denote the amount of bound streptavidin and biotinylated antibodies, respectively. Δf_EV,max_ is the maximum association observed for the given EV solution, Δf_EV,stable_ indicates the mass remaining after 20 minutes of dissociation. (c) CP tightly associates to the bare SiO_2_ surface of the QCM sensor chip (solid line; decrease in the frequency). Lipid vesicles bind (dashed line; initial frequency decrease) and subsequently fuse on the SiO_2_ surface over time thereby releasing their liquid content and forming stable supported lipid bilayers after lipid fusion (subsequent frequency increase).

**Figure 5.**
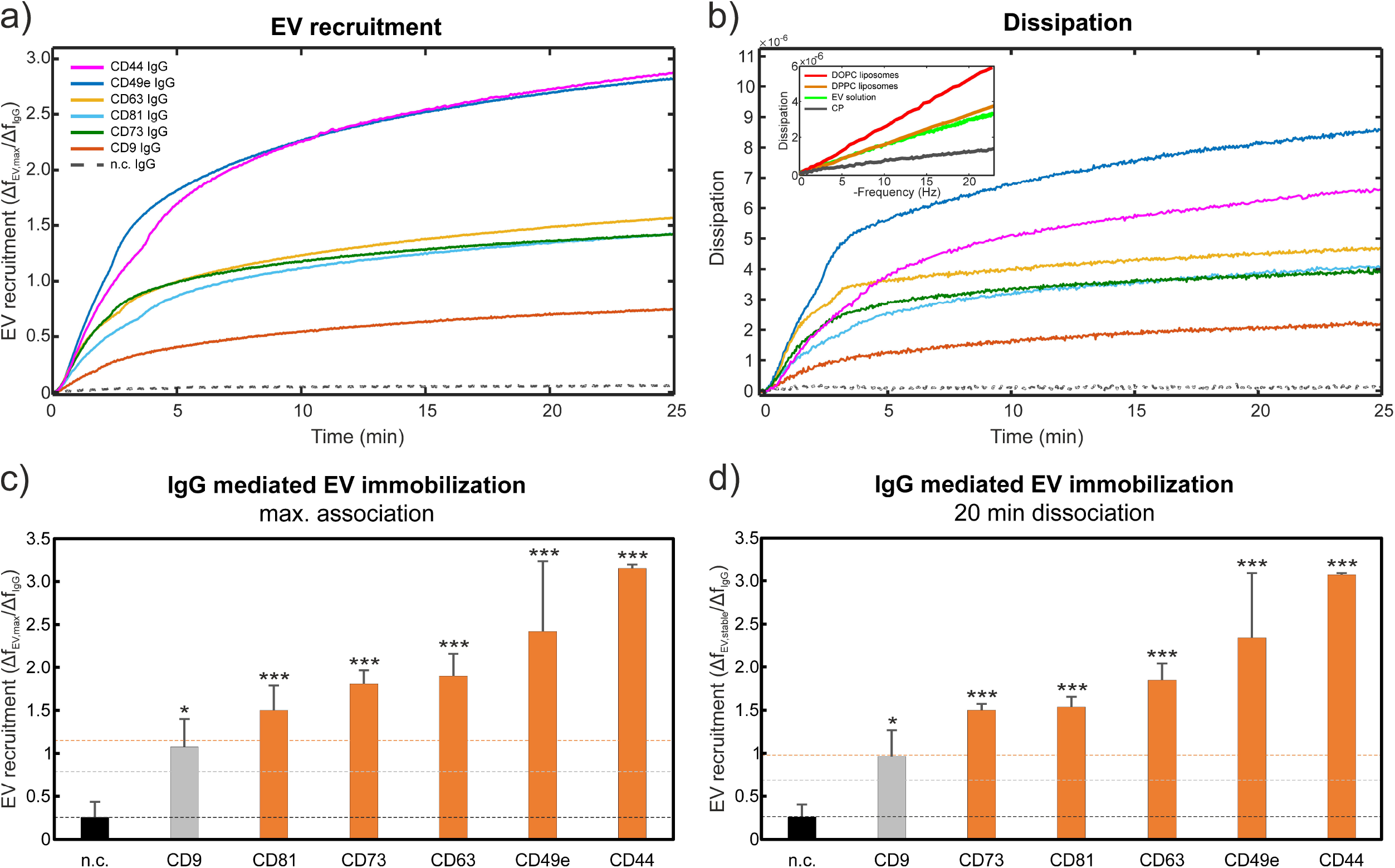
Analysis of EVs binding to IgG using QCM. (a) EV recruitment calculated from the frequency shift of EV binding normalized to the total number of immobilized CD9, CD44, CD49e, CD63, CD73 and CD81 antibodies for CD9, CD44, CD49e, CD63, CD73 and CD81 positive primary UC-MSC EVs and the negative control mouse IgG (dashed line). (b) Dissipation monitoring revealed an increase in dissipative energy loss after EV application. **Inset:** Correlation of frequency shift and dissipation signals obtained from applying either the EV solution, synthetic lipid vesicles (DOPC, DPPC), or CP to the QCM sensor chip. Dissipative properties of EV solutions closely resembles the properties of gel-phase lipid vesicles (DPPC) rather than more rigid CP. All data relates to the third harmonic frequency of the 5 MHz quartz crystals used. (c) and (d) Analysis of maximal (c) and final (d, after 20 min dissociation) relative EV recruitment levels of CD44, CD49e, CD61, CD73 and CD81 positive primary UC-MSC-EVs to the respective antibodies immobilized on the QCM chip. (d) Bars represent the mean of experimental replicates; error bars represent one standard deviation from the mean. The black dashed line represents the mean of all negative controls (n.c.). Grey and orange dashed lines represent the mean of all negative controls plus 3 and 5 standard deviations, respectively. All data relates to the third harmonic frequency of the 5 MHz quartz crystals used. Statistical analysis was performed using ordinary one-way Anova, Dunnett’s multiple comparisons test. *…p<0.05; **…p<0.01; ***…p<0.001, N=3-6.

## Discussion

Both EV based research and clinical applications have seen a surge in interest in recent years. Accordingly, experts in the field have established quality standards to ensure safety and efficacy for patients and enable efficient comparison of data between laboratories. When characterizing EV formulations it is critical to recognize that even with careful enrichment and purification steps a population of non-vesicle particles may be enriched alongside the EVs, as also stated in the MISEV 2018 guidelines. Separating experimental results stemming from EVs and NVPs may hence be an important prerequisite for the robust characterization of EV functionality, making it necessary to expand the current suit of standardized techniques. Here we present such an extended approach that provides data on EV quality and quantity using proven techniques and makes it possible to discriminate between EVs and NVPs. We first confirmed the presence of both populations using AFM and then used this complex mixture in QCM-I to selectively characterize EV surface proteins by carefully designing and monitoring the sensor properties. QI™-mode AFM imaging has been applied for quantification of the local elasticity of the EVs. Nevertheless, the more widely available tapping mode phase imaging can as well be used to image similar effects (Sharma et al., 2010). UC-or Wharton’s jelly-derived MSC have demonstrated a high safety and efficacy profile in more than 90 clinical studies, with therapeutic effects in different pathological conditions including neurological, hematological, immunological, liver, cardiac, endocrine, musculoskeletal, skin, ophthalmological and pulmonary diseases (Can et al., 2017). MSC-based immunomodulatory effects have mostly been attributed to the immunoregulatory properties of the MSC secretome including the contained membranous particle fraction of EVs, apoptotic bodies, microvesicles and exosomes (Harrell et al., 2019). Lai et al. investigated 3 different types of MSC EVs and could show that MSC produce at least 3 distinct 100 nm EV types, cholera toxin B chain (CTB)-, Annexin V (AV)- and Shiga toxin B subunit (ST)-EVs, that could be distinguished by their membrane lipid composition as well as their proteome and RNA cargo. They stated that CTB-binding EVs were CD81 positive and derived from endosomes and suggested that AV-EVs were derived from membrane organelles in the cytoplasm and ST-EVs have probably a biogenesis from the nucleus (Lai et al., 2016). To ensure reproducibility and quality of EV-based therapeutics, establishment of a standard and reliable quality control for the contained particle subtypes is important. For instance, ISEV suggests demonstrating the presence of three different kinds of proteins on EVs, including at least one transmembrane or lipid-bound extracellular protein (CD9, CD63, CD81), which have been originally established as exosome markers. Methods like ELISA, bead-based flow cytometry, aptamer- and carbon nanotube-based colorimetric assays, and SPR on surfaces such as antibody-coated nanorods can be used to quantify the amount of one or more specific molecules in the EV preparation (Witwer et al., 2019). In a recent proof-of-concept study, Suthar et al. (Suthar et al., 2020) used QCM-D to detect UC-MSC-EVs and monitored their binding to anti-CD63 antibodies with a frequency-based limit of detection of 2.9 × 10^8^ exosome sized particles/mL. This work suggests QCM-D as an attractive method for EV surface marker characterization, but the presented implementation using QCM-D, while highly sensitive, comes with a major barrier of entry in terms of instrumentation costs. Additionally, sample preparation methods based on self-assembled monolayers and covalently immobilized IgGs are challenging to establish, time-consuming, and limit sensor reusability. To address these issues, and to make QCM-based EV characterization accessible to a wider audience, we have established a label-free immunosensing method using QCM-I based on modular and fully reversible in-device functionalization of the sensor surface. Our approach represents a flexible (lipids are adaptable to the specific application, IgGs can be exchanged easily), reproducible, and economical (chips can be reused, sample consumption is minimized, inexpensive device compared to QCM-D) way to analyze EVs and their surface proteins. We have demonstrated the presence of all three EV-specific transmembrane proteins CD9, CD63, and CD81, thus corroborating MACSplex results and the statement of MISEV 2018 (Thery et al., 2018). In terms of functionality, UC-MSC-EVs express several adhesion molecules (CD29, CD44 and CD73), which facilitate their homing to the injured and inflamed tissues (Harrell et al., 2019). We were able to detect these typical MSC (CD44 and CD73) and UC-MSC (CD49e) markers for immobilizing UC-MSC-EVs as well. This is in line with our standard characterization protocols using the MACSplex analysis that reveal the presence of these surface markers in UC-MSC-EV preparations. CD73, a GPI- anchored 5’-ecto-nucleotidase that drives adenosine signaling, is of special interest here, as Angioni et al. found a novel anti-angiogenic mechanism with CD73-positive MSC-EVs responsible for adenosine-mediated modulation of angiogenesis in cancer (Angioni et al., 2020). CD73 positive MSC exosomes cause the phosphorylation of ERK1/2 and AKT kinases and elicited pro-survival signaling via these two latter enzymes (Lai et al., 2013). The conversion of AMP to adenosine by CD73 also triggers the immunomodulatory environment via T cells (Antonioli et al., 2019). Adenosine signaling via the A2a receptor is a costimulatory way in BDNF signaling and thereby intensifies the neuroprotective effect of BDNF (Pradhan et al., 2019). CD73 could play a dual role in mediating immunomodulation, survival and cytoprotection in the target tissue. Moreover, Müller et al. have shown that EV-bound CD73 in adipocyte-derived “adiposomes” (a mixture of exosomes or small EVs and larger-sized EVs) that is embedded in the outer leaflet of the plasma membrane via a GPI anchor can be transferred to the membranes of target cells (Muller et al., 2010). Thus, the enzymatic activity of CD73 can continue to generate adenosine-elicited signals via the target cell membrane. A prerequisite for this, however, is that CD73 is transferred as an intact enzyme that retains its capability of converting AMP to adenosine and phosphate. CD73 activity has been confirmed on MSC-derived plasma membrane vesicles (CDVs that are derived by forced extrusion of UC-MSCs) as well as on naturally secreted UC-MSC-derived EVs. The integrin α5, known as CD49e, was shown to promote osteogenic differentiation potential of UC-MSC (Zheng et al., 2018). Thus, we suggest that the association of these markers with functional membranous vesicles in addition to determining the MSC origin of EVs represents a valuable quality characteristic.

In conclusion, our combined approach, while not suited for high-throughput screening, enabled us to detect and characterize a suit of specific surface markers on membranous vesicles in a complex and clinically relevant EV formulation. QCM-I proved especially valuable to this end by not only providing the EV recruitment efficiency of a panel of EV markers but also simultaneously recording the viscoelastic properties of adsorbates, thus ensuring that all investigated markers immobilized EVs and not protein aggregates. Finally, our modular and reversible in-device sensor functionalization makes QCM it a highly promising and accessible technique for the specific detection of particle subpopulations.

## Author Contributions

EP, JS, JJ and MG made substantial contributions to conception and design, enquired and drafted the manuscript. EP, JS, BB, FW, CA, DA, EG and MN performed the scientific experiments and analyzed data. SW, JG, JP, JJ and MG have given final approval and revised the manuscript critically. All authors read and approved the final manuscript.

## Acknowledgements

We thank Alain Brisson for the cryo-EM images. This work was supported by the Austrian Forschungsförderungsgesellschaft Coin project “BioCETA” (No. 15379797), the European Fund for Regional Development (EFRE, IWB2020), and the Federal State of Upper Austria.

---

**Supplemental Figure 1.** Exemplary AFM topography and elasticity image of EVs and NVPs. (a) and (b) Overview image of three EVs and three NVPs marked exemplarily with a white or green circle and (c) and (d) exemplary AFM image of four EVs of various sizes. The corresponding elasticity-profiles of the marked particles displayed below show the typical W-profile (“doughnut shape”) to varying degrees.

