## Supplementary figures and images for "Comprehensive label-free characterization of extracellular vesicles and their surface proteins"

### Supplemental Figure 1

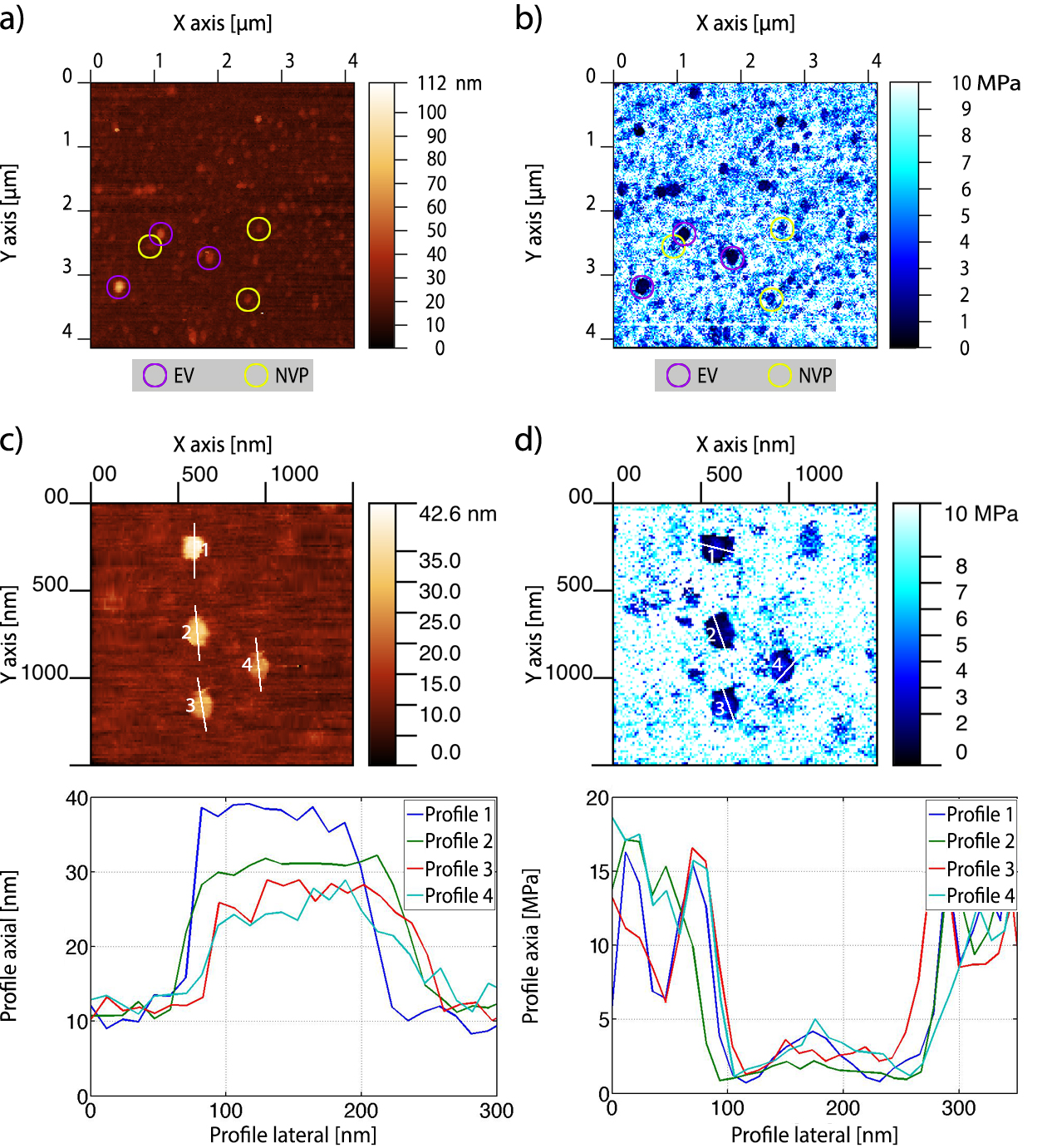
